# TeachEnG: a <u>Teach</u>ing <u>En</u>gine for <u>G</u>enomics

**DOI:** 10.1101/111054

**Authors:** Minji Kim, Yeonsung Kim, Lei Qian, Jun S. Song

**Affiliations:** Department of Electrical and Computer Engineering, University of Illinois at Urbana-Champaign, Urbana, IL 61801, USA; Carl R. Woese Institute for Genomic Biology, University of Illinois at Urbana-Champaign, Urbana, IL 61801, USA; Department of Mathematics and Computer Science, Fisk University, Nashville, TN 37208; Department of Bioengineering, University of Illinois at Urbana-Champaign, Urbana, IL 61801, USA; Department of Physics, University of Illinois at Urbana-Champaign, Urbana, IL 61801, USA

## Abstract

**Motivation:** Bioinformatics is a rapidly growing field that has emerged from the synergy of computer science, statistics, and biology. Given the interdisciplinary nature of bioinformatics, many students from diverse fields struggle with grasping bioinformatic concepts only from classroom lectures. Interactive tools for helping students reinforce their learning would be thus desirable. Here, we present an interactive online educational tool called TeachEnG (acronym for <u>Teach</u>ing <u>En</u>gine for <u>G</u>enomics) for reinforcing key concepts in sequence alignment and phylogenetic tree reconstruction. Our instructional games allow students to align sequences by hand, fill out the dynamic programming matrix in the Needleman-Wunsch global sequence alignment algorithm, and reconstruct phylogenetic trees via the maximum parsimony and Unweighted Pair Group Method with Arithmetic mean (UPGMA) algorithms. With an easily accessible interface and instant visual feedback, TeachEnG will help promote active learning in bioinformatics.

**Availability and Implementation:** TeachEnG is freely available at http://song.igb.illinois.edu/TeachEnG/. It is written in JavaScript and compatible with Firefox, Safari, Chrome, and Microsoft Edge.

## 1 Introduction

Many of the tools in bioinformatics are based on algorithms developed in mathematics, statistics, and computer science. For instance, sequence alignment and RNA folding prediction algorithms are based on dynamic programming (Bellman, 1952). Popular examples include the Needle-man-Wunsch (NW) global sequence alignment algorithm (Needleman and Wunsch, 1970) and the Smith-Waterman local alignment algorithm. Another important concept frequently taught in bioinformatics courses is phylogenetic tree reconstruction via the maximum parsimony (Fitch, 1971) and the Unweighted Pair Group Method with Arithmetic mean (UPGMA) (Sokal and Michener, 1958) algorithms. Given the interdisciplinary nature of bioinformatics, understanding these algorithms and mathematical concepts often pose challenges to students from diverse fields, including experimental biology. Hands-on training to reinforce learning would be ideal, but such opportunities are not readily available, especially in small teaching colleges. The developers of SAT (Ibarra and Melo, 2010) attempt to address this problem by providing a java package that fills out the dynamic programming matrix for sequence alignment, but the users do not have the option to manually fill out their own matrices and receive feedback. Furthermore, to the best of our knowledge, there is no software tool for teaching phylogenetic tree reconstruction.

The National Institutes of Health (NIH) has recently funded Big Data to Knowledge (BD2K) Centers and various BD2K Training projects to tackle this type of challenges and to broadly disseminate educational computational tools. As part of the KnowEnG BD2K Center, we have developed a web-based resource called TeachEnG (acronym for <u>Teach</u>ing <u>En</u>gine for <u>G</u>enomics, and pronounced “teaching”) to help students reinforce their understanding of algorithms widely used in bioinformatics. Our website currently contains four different modules that are implemented as interactive games. The “Sequence Alignment” game asks the user to rearrange two sequences to get as many matches and as few mismatches and gaps as possible, and demonstrates the need for a constructive sequence alignment algorithm. In the subsequent portal “Sequence Alignment: Needleman-Wunsch”, the user is given two sequences and penalties for matches, mismatches, and gaps. The user then fills out the dynamic programming matrix in the NW algorithm. A natural extension to sequence alignment is to compare multiple sequences from more than two species and find evolutionary trees. Our last two modules “Maximum Parsimony” and “UPGMA” allow the user to reconstruct phylogenetic trees from a given set of sequences and similarity measures for four species.

## 2 Software Description

### 2.1 Sequence Alignment

The goal of sequence alignment is to arrange two sequences in the most similar manner. Here, the similarity is measured by a weighted sum of matches, mismatches, and gaps. In the Sequence Alignment game, consisting of three levels of difficulty, the user is prompted to manually arrange two sequences to achieve the highest score. This time-consuming exercise highlights the need for a fast algorithm to complete the task, leading us to the NW algorithm.

In the NW game, the user is given two sequences **v** and **w** of lengths *n* and *m*, respectively, and a scoring matrix δ. The goal is to fill out the dynamic programming matrix S in the NW algorithm. For each entry S_i,j_ in the (*n*+l)×(*m*+l) matrix, the user needs to put correctly the maximum of three values: S_i−1,j−1_+ δ(v_i_,w_j_) (match/mismatch), S_i−1_,_j_+δ(Vi,-) (gap in **w**), and S_i,j−1_+ δ(-,Wj) (gap in **v**). The program automatically checks the user input at every step and informs the user. After the matrix is completed, the program will backtrack the paths and return the best alignment path. Fig. 1A shows a screenshot of the NW game in progress.

**Fig. 1.**
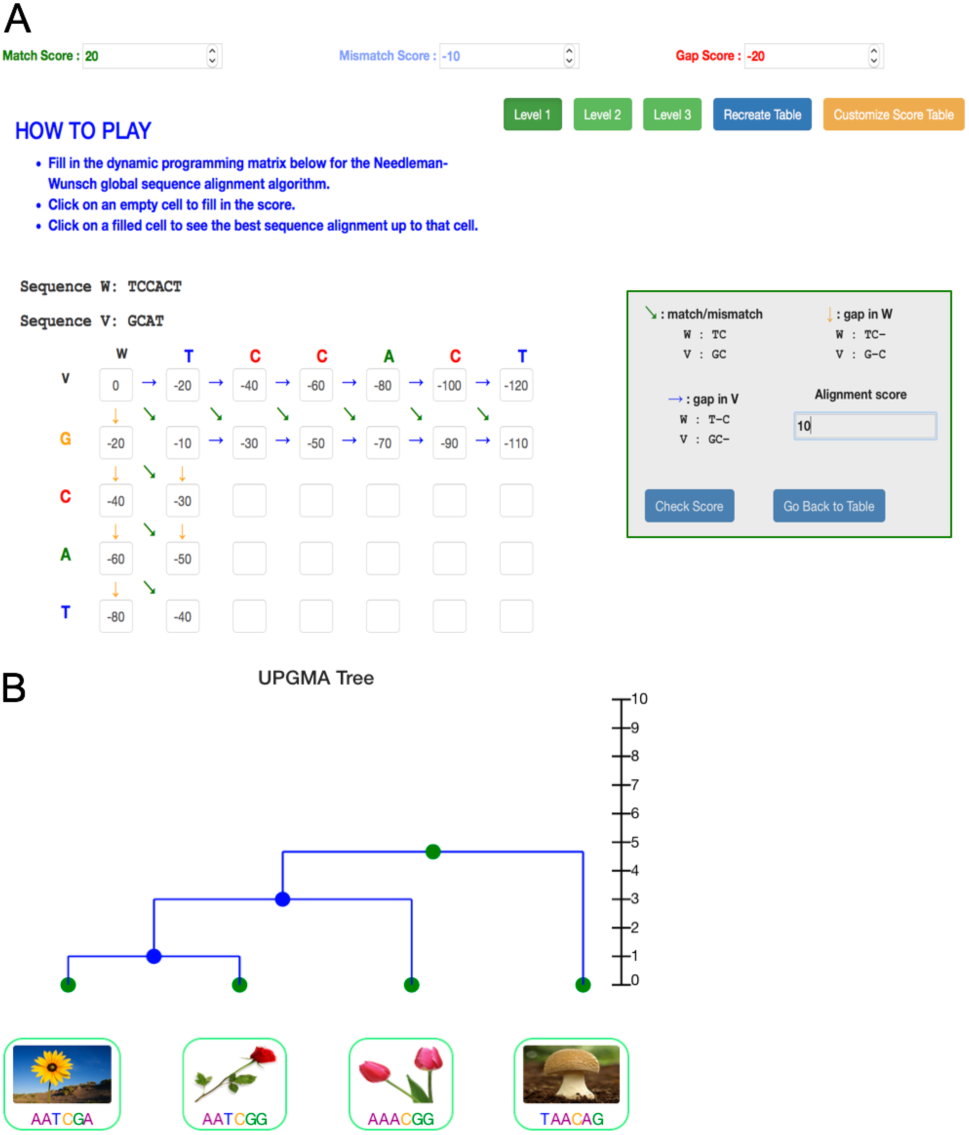
Snapshots of TeachEnG. (A) An example of the dynamic programming matrix for the Needleman-Wunsch global sequence alignment algorithm and TeachEnG’s user interface box. (B) An example of final UPGMA phylogenetic tree.

In both games, the score matrix can be customized by the user.

### 2.2 Phylogenetic Tree Reconstruction

Sequence similarity explored in the previous section can be extended to comparing sequences from multiple species and reconstructing phylogenetic trees to represent evolutionary relationships. One popular method is to impose maximum parsimony and is implemented in our third module. Given a set of four species and their corresponding short sequences, the user first guesses the correct tree by grouping similar species together; the guessed tree will be compared with the maximum parsimony tree at the final stage of the game. The user then has to select the correct informative sites to proceed to the next stage of the game and subsequently interacts with the game to evaluate the three possible tree structures. The final webpage selects the tree with the minimum score as the maximum parsimony tree and compares it to the user’s initial guess.

Another popular method is based on a pre-defined distance measure between species, where short distance implies high similarity. In our last module, we implemented the UPGMA algorithm, where the user is given a distance matrix, and the goal is to iteratively select the pair with the shortest distance, merge the species, and then fill out the new distance matrix for merged nodes. When all species are grouped into one common ancestor, the program returns the completed UPGMA tree (Fig. 1B).

## 3 Conclusion

We developed an interactive tool called TeachEnG for teaching widely used algorithms in bioinformatics. The user can manually align two sequences, fill out the dynamic programming matrix, and reconstruct evolutionary trees. This interactive resource will complement lecture-based teaching methods by providing ample practice problems and instant visual feedback. TeachEnG is expandable and can easily incorporate new educational games in the future. Fisk University is an NIH-supported partner of the KnowEnG Center and is currently using TeachEnG in their bioinformatics course. We hope that TeachEnG will help advance the broader bioinformatics community by educating students and researchers from diverse backgrounds.

## Acknowledgements

The authors would like to thank Jiaxin Liu for helpful suggestions.

## References

Bellman, R. (1952.) On the theory of dynamic programming. P. Natl. Acad. Sci., 38(8), 716–719.

Fitch, W.M. (1971.) Toward defining the course of evolution: minimum change for a specific tree topology. Syst. Biol., 20(4), 406–416.

Ibarra, I.L. and Melo, F. (2010). Interactive software tool to comprehend the calculation of optimal sequence alignments with dynamic programming. Bioinformatics, 26(13), 1664–1665.

Needleman, S.B. and Wunsch, C.D. (1970). A general method applicable to the search for similarities in the amino acid sequence of two proteins. J. Mol. Biol., 48(3), 443–453.

Sokal, R.R. and Michener, C.D. (1958). A statistical method for evaluating systematic relationships. Univ. Kans. Sci. Bull., 38, 1409–1438.

